# Near-complete genome assembly of tomato (*Solanum lycopersicum*) cultivar Micro-Tom

**DOI:** 10.1101/2023.10.26.564283

**Authors:** Kenta Shirasawa, Tohru Ariizumi

**Author notes:** This work is dedicated to Tohru Ariizumi, who passed away on May 10, 2023. To whom correspondence should be addressed: Kenta Shirasawa.

## Abstract

We present a near-complete genome assembly of tomato (*Solanum lycopersicum*) cultivar Micro-Tom, which has been recognized as a model cultivar for fruit research. The genome DNA of Micro-Tom, provided by the National BioResource Project (NBRP) Tomato of Japan, was sequenced to obtain 72 Gb of high-fidelity long reads. These reads were assembled into 140 contigs, spanning 832.8 Mb, with an N50 length of 39.6 Mb. The contigs were aligned against the tomato reference genome sequence SL4.0 to establish a chromosome-level assembly. The genome assembly of Micro-Tom contained 98.5% complete BUSCOs and a total of 31,429 genes. Comparative genome structure analysis revealed that Micro-Tom possesses a cluster of rDNA genes spanning a 15 Mb stretch at the short arm of chromosome 2. This region was not found in the genome assemblies of previously sequenced tomato cultivars, possibly because of the inability of previous technologies to sequence such repetitive DNA. In conclusion, the near-complete genome assembly of Micro-Tom reported in this study would advance the genomics and genetics research on tomato and facilitate the breeding of improved tomato cultivars.

## Introduction

Tomato (*Solanum lycopersicum*) is a member of the Solanaceae family, which includes several plant species of agricultural and ornamental importance such as potato (*Solanum tuberosum*), eggplant (*Solanum melongena*), pepper (*Capsicum annuum*), tobacco (*Nicotiana tabacum*), and petunia (*Petunia* × *hybrida*). Because tomato has a diploid genome (2n = 2x =24) of small size (∼900 Mb), along with high consumption and production rates worldwide, its molecular biology has been studied since the 1980s (Shirasawa and Hirakawa 2013). Consequently, tomato genes controlling agronomically important traits have been identified and used in breeding programs, which has resulted in the development of functional genomics tools as well as experimental lines (Shirasawa and Hirakawa 2013).

Micro-Tom is a miniature tomato cultivar (Scott and Harbaugh 1989). Because of its compact plant size and short life cycle under laboratory conditions, Micro-Tom is considered a model tomato cultivar for research (Meissner et al. 1997). To accelerate research on the molecular biology of tomato, numerous genomics and genetic resources of Micro-Tom as well as artificial induced mutant lines have been developed so far (Saito et al. 2011; Carvalho et al. 2011; Just et al. 2013; Shikata et al. 2016). The National BioResource Project Tomato (NBRP Tomato) of Japan has collected, propagated, maintained, and distributed the Micro-Tom bioresources to the research community, with the aim to promote functional genomics studies in tomato (Ariizumi et al. 2011). Molecular techniques for screening mutated genes have been developed (Okabe et al. 2011), and full-length cDNA libraries have been constructed and sequenced (Aoki et al. 2010). A bacterial artificial chromosome (BAC) library has been also constructed, and the BAC-end sequences have been mapped on to the tomato genome (Asamizu et al. 2012). Genetic maps for Micro-Tom have been established for genetic studies (Shirasawa et al. 2010, 2016a). Owing to the publication of the whole-genome sequence of tomato cultivar Heinz 1706 (Tomato Genome Consortium 2012), sequence variants in Micro-Tom mutants could be identified using a whole-genome resequencing strategy (Shirasawa et al. 2016b). In parallel, spontaneous polymorphisms within wild-type Micro-Tom lines were also found (Shirasawa et al. 2010, 2016b; Kobayashi et al. 2014), suggesting that Micro-Tom consists of multiple lines, which could be divided into at least four genetically distinguishable groups (i.e., France, USA, NBRP-Japan, and NIVTS-Japan) (Shirasawa et al. 2016b).

The genome of three lines of Micro-Tom has been sequenced to date. The first genome sequence of Micro-Tom was released by the National Polytechnic Institute of Toulouse, France, in 2020 (genome assembly ID, SLYMIC; GenBank accession number: JAAXDC000000000). This dataset comprises 12 chromosome sequences spanning a physical distance of 812.5 Mb, with undetermined sequences (i.e., gaps) spanning 22.6 Mb. The second genome sequence of Micro-Tom, reported by a Chinese group (microTom, http://eplant.njau.edu.cn/microTomBase) (Xue et al. 2023), represents a 798.9 Mb chromosome-level assembly containing only 44.7 kb gaps. The Japanses line of Micro-Tom, which has been maintained in Kazusa DNA Research Institute, Japan, was sequenced most recently (SLM_r1.2, GenBank accession number: BSVZ00000000) (Nagasaki et al. 2023), and represents a genome assembly of size 795.7 Mb, with a gap size of 14.0 Mb. These three assemblies were generated using short-read (Illumina, San Diego, CA, USA), long-read (PacBio, Menlo Park, CA, USA; Oxford Nanopore Technologies, Oxford, UK), linked-read (10X Genomics, Pleasanton, CA, USA), and/or optical mapping technologies (Bionano Genomics, San Diego, CA, USA). The currently available high-fidelity (HiFi) long-read sequencing technology of PacBio could be used to establish telomere-to-telomere genome assemblies (Kurokochi et al. 2023; Sato et al. 2023), which are continuous assemblies ranging from one end of a chromosome to the other end without gaps.

In this study, we employed the HiFi long-read sequencing technology to establish a high-quality (i.e., high genome coverage and long contiguity) chromosome-level genome assembly for Micro-Tom NBRP-Japan line, since most of the genomics and genetic resources of Micro-Tom are based on the NBRP-Japan line. The chromosome-level genome assembly of Micro-Tom generated in this study is the highest quality assembly reported to date. This genome assembly could be used to advance the functional genomics of tomato and to perform mutant screening in Micro-Tom.

## Materials and methods

### Plant materials

The NBRP-Japan line of Micro-Tom (TOMJPF00001) was used for whole-genome sequencing in this study. Additionally, mutations were detected in the genome of nine EMS-induced Micro-Tom mutants (TOMJPE2703, TOMJPE3484, TOMJPE5212-5, TOMJPE5406, TOMJPE5409, TOMJPE5770-1, TOMJPE5906, TOMJPW0604, and TOMJPW1559-1), which set large fruits, according to the TOMATOMA database (Saito et al. 2011; Shikata et al. 2016).

### Genome sequencing and assembly

Genome DNA was extracted from the young leaves of Micro-Tom lines using Genomic Tip (Qiagen, Hilden, Germany). The extracted DNA was sheared into 30 kb fragments using Megaruptor 2 (Diagenode, Seraing, Belgium), and then subjected to HiFi SMRTbell library preparation with the SMRTbell Express Template Prep Kit 2.0 (PacBio). The resultant library was separated on BluePippin (Sage Science, Beverly, MA, USA) to remove short DNA fragments (<15 kb), and sequenced with SMRT Cell 8 M on the Sequel II system (PacBio). The obtained HiFi reads were assembled with Hifiasm (Cheng et al. 2021) using default parameters. Potential contaminants, i.e., organellar and fungal DNA sequences, were identified based on a sequence similarity search in the UniProtKB database (UniProt Consortium 2023) using DIAMOND (Buchfink et al. 2021), with an E-value cutoff of <1E-10. The assembled sequences were aligned against the tomato reference genome sequence SL4.0 (https://solgenomics.net) using Ragoo (Alonge et al. 2019) to build pseudomolecule sequences. Telomere sequences containing repeats of a 7 bp motif (5’ -TTTAGGG-3’) were searched by the search subcommand of tidk (https://github.com/tolkit/telomeric-identifier), with a window size of 100,000 bp. The assembly quality was assessed with BUSCO (Simão et al. 2015).

### Prediction of genes and repeat sequences

Gene prediction was performed with BRAKER3 (Gabriel et al. 2023) based on the peptide sequences of the predicted genes of ITAG4.0 (https://solgenomics.net) and SLM_r1.2 (Nagasaki et al. 2023), full-length cDNA sequences of Micro-Tom (GenBank accession nos.: AB211519– AB211522, AB211526, AK224591–AK224910, AK246135–AK248077, and AK319176– AK330134), and RNA-Seq reads (NCBI Sequence Read Archive accession nos.: SRR12560324– SRR12560335) (Bae et al. 2021). Then, gene sequences reported in the genome assemblies of ITAG4.0 and SLM_r1.2 were mapped onto the pseudomolecule sequences using Liftoff (Shumate and Salzberg 2020). The genome positions of predicted and mapped genes were compared using the intersect command of BEDtools (Quinlan and Hall 2010). The predicted genes were functionally annotated using emapper implemented in EggNOG (Cantalapiedra et al. 2021), in conjunction with DIAMOND (Buchfink et al. 2021), against the UniProtKB database (UniProt Consortium 2023).

Repetitive sequences in the assembly were identified with RepeatMasker (https://www.repeatmasker.org) using repeat sequences registered in Repbase and a *de novo* repeat library built with RepeatModeler (https://www.repeatmasker.org).

### Comparative analysis of genome structures

The pseudomolecule sequences were aligned against SLYMIC, microTom, SLM_r1.2, and SL4.0 genome assemblies as references with UniMAP (https://github.com/lh3/unimap), and the alignments were visualized with D-Genies (Cabanettes and Klopp 2018).

### Whole-genome sequencing analysis of the Micro-Tom mutants

Genome DNA was extracted from the young leaves of Micro-Tom mutants using DNeasy Plant Mini Kit (Qiagen), and subjected to library preparation with the SPRIworks System I for the Illumina sequencer (Beckman Coulter, Brea, CA, USA). The nucleotide sequences of the resultant libraries were determined using HiSeq 1000 (Illumina) in paired-end, 101 bp mode. After removing low-quality bases (quality value < 10) with PRINSEQ (Schmieder and Edwards 2011) and adaptor sequences (AGATCGGAAGAGC) with fastx_clipper in the FASTX-Toolkit (http://hannonlab.cshl.edu/fastx_toolkit), the reads were mapped onto the pseudomolecule sequences with Bowtie2 (Langmead and Salzberg 2012). Sequence variants were detected using the mpileup and call commands of BCFtools (Li 2011), and high-confidence biallelic SNPs were identified with VCFtools (Danecek et al. 2011) using the following parameters: minimum read depth ≥ 8 (--minDP 8); minimum variant quality = 20 (--minQ 20); maximum missing data < 0.5 (--max-missing 0.5); and minor allele frequency ≥ 0.05 (--maf 0.05). Effects of SNPs on gene function were estimated with SnpEff (Cingolani et al. 2012).

## Results

### Genome assembly

We obtained a total of 72.3 Gb HiFi data, consisting of 1.9 M reads (N50 = 26 kb), from two SMRT Cells (8M). The reads were assembled into 545 contigs (N50 = 39.6 Mb) spanning a physical distance of 866.5 Mb. Potentially contaminating sequences derived from organellar and fungal DNA were removed to obtain 140 contigs, totaling 832.8 Mb with an N50 value of 39.6 Mb (Table 1). This assembly was designated as SLM_r2.0. All 140 contigs were aligned against the standard tomato reference genome sequence SL4.0, and 12 chromosome-scale pseudomolecule sequences were established (SLM_r2.0.pmol; Tables 1 and 2, Figure 1). The complete BUSCO score for SLM_r2.0.pmol was 98.5% (Table 3). Telomeric repeats were found at both ends of four chromosomes (chr. 3, 7, 8, and 9) and at one end of the remaining eight chromosomes (chr. 1, 2, 4, 5, 6, 10, 11, and 12).

**Table 1.**
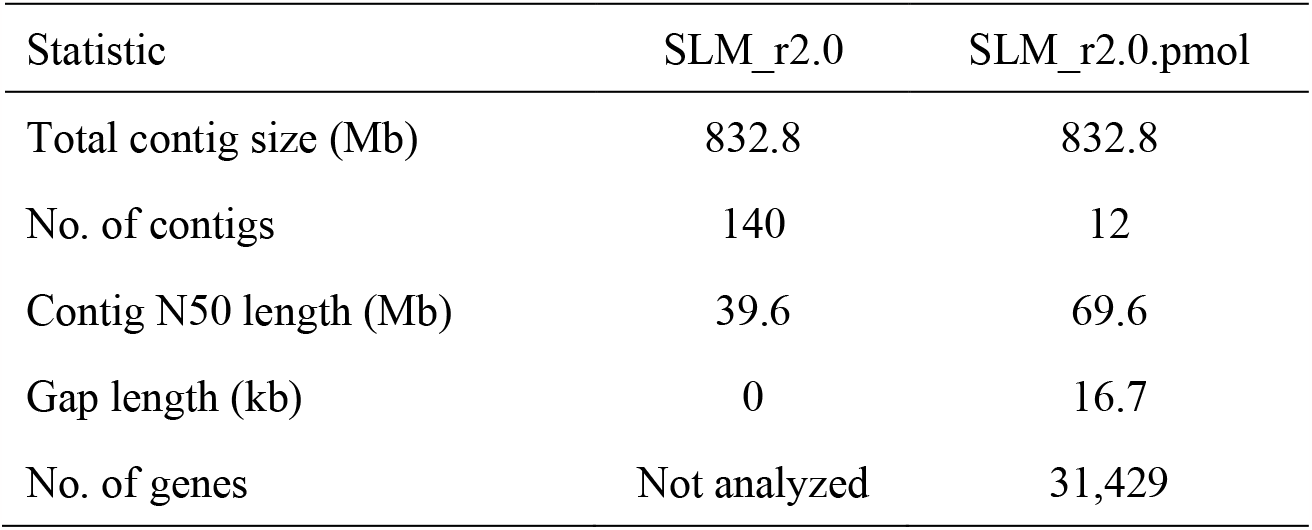
Statistics of the genome assembly of Micro-Tom NBRP-Japan line.

**Figure 1.**
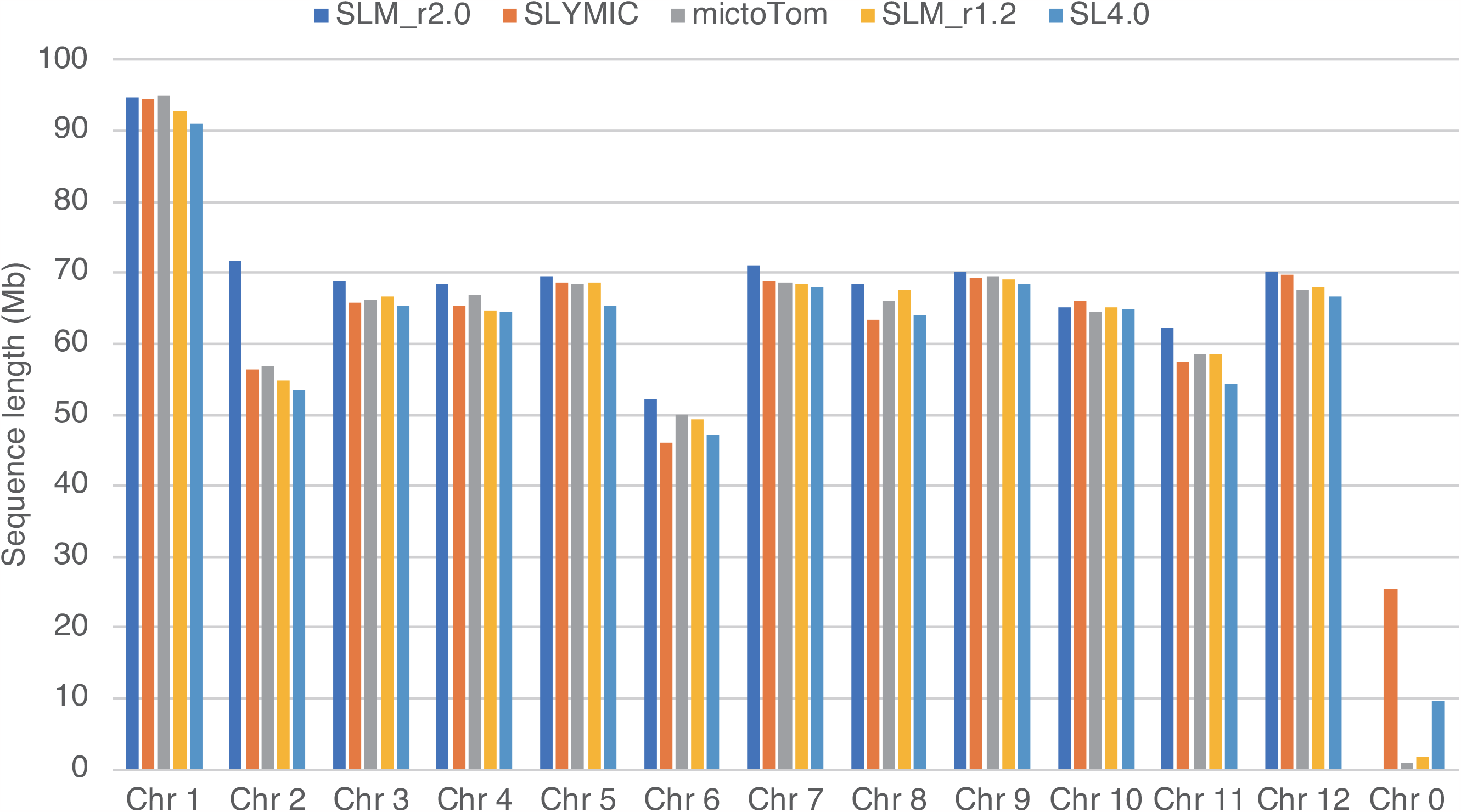
Sequence lengths of chromosome-level genome assemblies of tomato. Chr 1 to Chr 12 represent chromosome numbers 1 to 12 in tomato. Chr 0 indicates sequences unassigned to any chromosomes.

### Gene and repeat prediction

A total of 45,868 protein-coding genes were initially predicted in the SLM_r2.0.pmol assembly. After removing potential transposable elements, 31,429 genes were selected as high-confidence (HC) genes (Table 2), while the remaining 14,439 genes were designated as low-confidence (LC) genes. The complete BUSCO score for the HC genes was 93.9% (Table 3). In parallel, 34,451 SLM_r1.2 and 33,823 SL4.0 gene models were mapped onto SLM_r2.0.pmol. The genomic positions of 25,706 out of 31,429 HC genes were overlapped with 24,626 SLM_r1.2 and/or 24,526 SL4.0 genes. A total of 43,753 genes were represented in at least one of the three genome assemblies.

**Table 2.** Statistics of Micro-Tom pseudomolecule sequences (SLM_r2.0.pmol)

| Chromosome | Total length |  | Number of genes | Percentage of genes<br>(relative to genes in the<br>entire genome) |
| --- | --- | --- | --- | --- |
|  | (bp) | (%) |  |  |
| SLM_r2.0ch01 | 94,708,454 | 11.4 | 3,439 | 10.9 |
| SLM_r2.0ch02 | 71,710,245 | 8.6 | 3,343 | 10.6 |
| SLM_r2.0ch03 | 68,736,248 | 8.3 | 2,981 | 9.5 |
| SLM_r2.0ch04 | 68,494,551 | 8.2 | 2,851 | 9.1 |
| SLM_r2.0ch05 | 69,564,429 | 8.4 | 1,980 | 6.3 |
| SLM_r2.0ch06 | 52,172,941 | 6.3 | 2,683 | 8.5 |
| SLM_r2.0ch07 | 71,053,079 | 8.5 | 3,120 | 9.9 |
| SLM_r2.0ch08 | 68,473,177 | 8.2 | 2,483 | 7.9 |
| SLM_r2.0ch09 | 70,139,957 | 8.4 | 2,248 | 7.2 |
| SLM_r2.0ch10 | 65,184,273 | 7.8 | 2,014 | 6.4 |
| SLM_r2.0ch11 | 62,371,966 | 7.5 | 2,212 | 7.0 |
| SLM_r2.0ch12 | 70,164,003 | 8.4 | 2,075 | 6.6 |
| Total | 832,773,323 | 100.0 | 31,429 | 100.0 |

**Table 3.** Completeness evaluation of genome assembly and predicted genes.

| BUSCO type | Genome (SLM_r2.0.pmol) | Predicted genes |
| --- | --- | --- |
| Complete | 98.5% | 93.9% |
| Single-copy | 97.8% | 93.2% |
| Duplicated | 0.7% | 0.7% |
| Fragmented | 0.4% | 1.1% |
| Missing | 1.1% | 5.0% |

Repetitive sequences occupied a total physical distance of 610.8 Mb (73.3%) in the SLM_r2.0.pmol genome assembly. Nine major types of repeats were identified in varying proportions (Table 4). The dominant repeat types in the chromosome sequences were long-terminal repeats (36.4%, 303.5 Mb) including *Gypsy*-(29.2%, 243.1 Mb) and *Copia*-type (6.3%, 52.5 Mb) retroelements. Repeat sequences unavailable in public databases totaled 238.1 Mb (28.6%).

**Table 4.** Repetitive sequences in the Micro-Tom chromosome-level genome assembly.

(SLM\_r2.0.pmol)
| Repeat type | No. of elements | Length occupied (bp) | Percentage of<br>repetitive sequences<br>(relative to length of<br>the entire genome) |
| --- | --- | --- | --- |
| SINEs | 6,313 | 789,544 | 0.09 |
| LINEs | 34,374 | 14,573,305 | 1.75 |
| LTR elements | 210,540 | 303,544,452 | 36.43 |
| DNA transposons | 98,905 | 30,015,504 | 3.6 |
| Small RNA | 9,167 | 13,323,116 | 1.6 |
| Satellites | 2,244 | 808,584 | 0.1 |
| Simple repeats | 105,416 | 7,238,810 | 0.87 |
| Low complexity repeats | 21,347 | 1,078,936 | 0.13 |
| Unclassified | 545,763 | 238,127,988 | 28.58 |

### Comparative analysis of genome structures

The SLM_r2.0.pmol genome assembly covered the entire genome of four tomato lines, i.e., SLYMIC, microTom, SLM_r1.2, and SL4.0 (Figure 2). However, potential structural variations were found between SLM_r2.0.pmol and the above-mentioned four lines. The most prominent difference, approximately 15 Mb in length, was found at the short arm of the chromosome 2. This nucleotide sequence was present in SLM_r2.0.pmol but was absent from SLYMIC, microTom, SLM_r1.2, and SL4.0. The top of tomato chromosome 2 has been reported to contain highly repetitive rDNA sequences (Vallejos et al. 1986). Indeed, in accordance with gene annotations, a total of 588 genes, including 259 copies of genes similar to the regulator of rDNA transcription protein 15 (UniProt accession no.: A0A6N2C889) and 222 copies of uncharacterized proteins (UniProt accession no.: A0A2G2UY24), were repetitively found at the top of chromosome 2 in SLM_r2.0.pmol.

**Figure 2.**
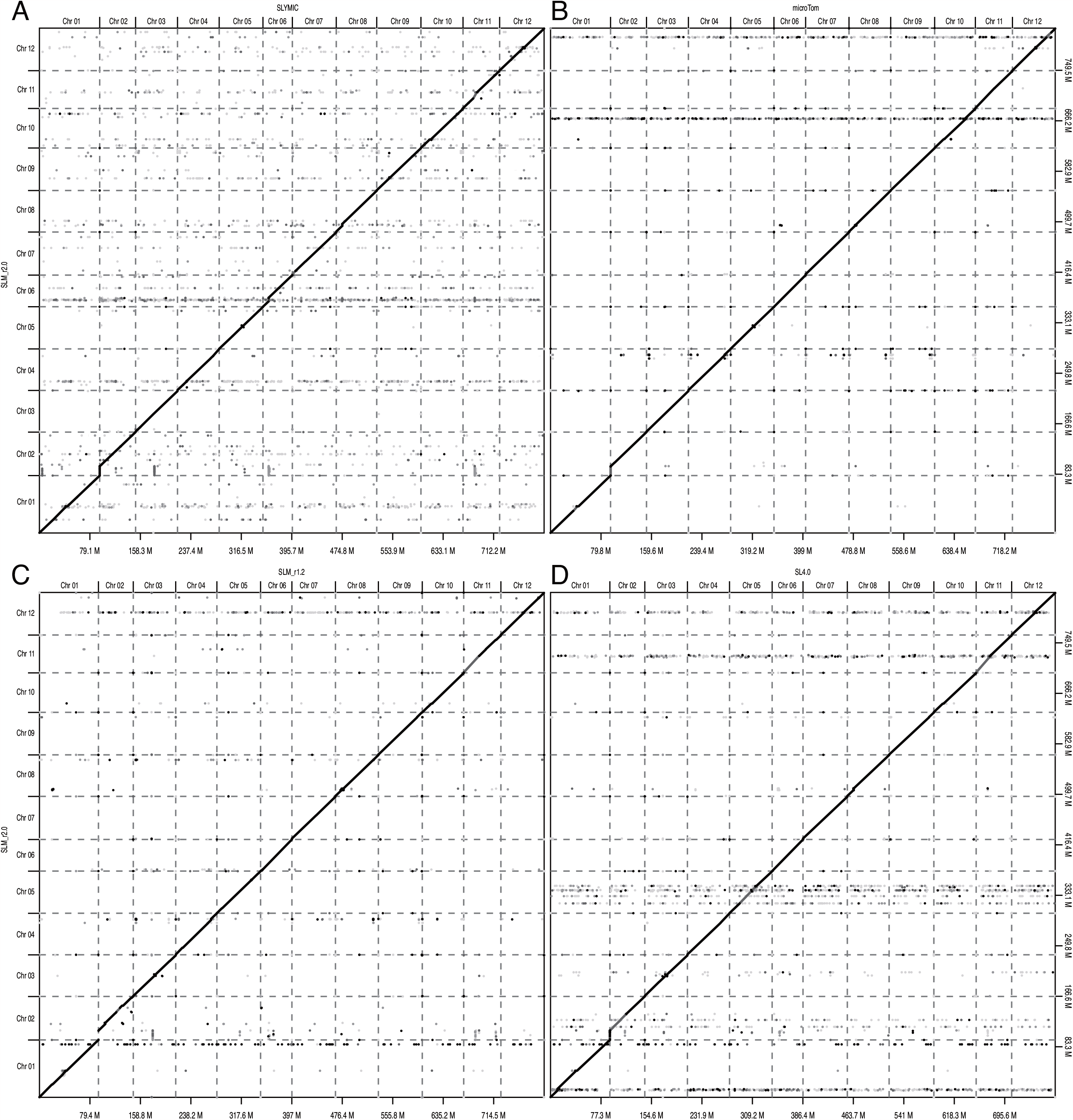
Comparative analysis of the genome sequence and structure of Micro-Tom lines, with Heinz 1706 as the standard line. (A–D) Plots showing SLYMIC (A), microTom (B), SLM_r1.2 (C), and SL4.0 (D) assemblies on the x-axes, with the SLM_r2.0.pmol assembly on the y-axes. Chromosome names are indicated above the x-axis and on the left side of the y-axis, and genome sizes (Mb) are shown below the x-axis and on the right side of the y-axis.

### Genome analysis of large-fruit mutants

A total of 265.3 Gb paired-end reads were obtained from nine large-fruit mutants of Micro-Tom. Paired-end reads of four wild-type tomato lines downloaded from a public DNA database (France: ERR340383 and ERR340384; USA: DRR118571; NBRP-Japan: DRR000741; and NIVTS-Japan: DRA002470) were included as controls. The reads were mapped on to SLM_r2.0.pmol, and a total of 172,791 sequence variants were identified. Of the 172,791 sequence variants, 8,601 variants uniquely found in a single mutant line were selected as induced mutations, whereas the other 164,190 variants were spontaneous sequence polymorphisms shared by multiple lines. On average, one induced mutation was found every 0.87 Mb distance (= 8,601 induced mutations / 832.8 Mb genome × 9 mutants). The four wild-type lines were genetically distinguishable, and all nine mutants were classified in the NBRP-Japan group (as expected), except one (TOMJPE2703), which was classified in the NIVTS-Japan group. The 8,601 induced mutations included 56 indels and 8,545 single nucleotide variations, of which 6,501 (76.1%) were G/C to A/T transitions. Out of 8,601 induced mutations, 693 mutations (8.1%) were located within genes, whereas the other 7,908 mutations (91.9%) were found outside of genic regions. Among the induced mutations, 14 (0.2%) were deleterious mutations, such as frameshift mutations, missense mutations at the first codon, nonsense mutations, and splice donor and acceptor variants, which disrupt gene function. Therefore, 14 genes harboring deleterious mutations were selected as candidates likely responsible for the large-fruit phenotype of the nine mutants (Table 5).

**Table 5.** Genes with deleterious mutations in large-fruit Micro-Tom mutants.

| Gene ID | Mutation | Mutant line | Gene annotation |
| --- | --- | --- | --- |
| SLM2ch01g03908 | Missense | TOMJPE5906 | MADS-box transcription factor |
| SLM2ch02g07037 | Nonsense | TOMJPE5409 | Mate efflux family protein |
| SLM2ch03g10021 | Splice donor variant | TOMJPE5212-5 | Amino-acid N-acetyltransferase |
| SLM2ch03g13085 | Nonsense | TOMJPE5770-1 | NMDA receptor-regulated protein 1 |
| SLM2ch04g13789 | Splice donor variant | TOMJPE3484 | PHD domain-containing protein |
| SLM2ch06g23428 | Nonsense | TOMJPE5906 | Transcriptional regulator of RNA polII, SAGA, subunit |
| SLM2ch07g27934 | Nonsense | TOMJPE5770-1 | Belongs to the short-chain dehydrogenases reductases family |
| SLM2ch08g31097 | Missense | TOMJPE5906 | Histidine decarboxylase |
| SLM2ch09g32719 | Nonsense | TOMJPE5212-5 | Splicing factor 3A subunit |
| SLM2ch09g35243 | Splice acceptor variant | TOMJPW0604 | Serine threonine-protein phosphatase |
| SLM2ch10g36090 | Splice acceptor variant | TOMJPE5906 | Purple acid phosphatase |
| SLM2ch12g42744 | Frameshift | TOMJPW0604 | Uncharacterized protein |
| SLM2ch12g42984 | Nonsense | TOMJPE5212-5 | Histone H2B |
| SLM2ch12g44447 | Splice acceptor variant | TOMJPW0604 | Uncharacterized protein |

## Discussion

We present a near-complete chromosome-scale genome assembly of the tomato cultivar Micro-Tom (SLM_r2.0.pmol), which spanned 832.8 Mb in length (Tables 1 and 2). The length of SLM_r2.0.pmol assembly was greater than those of Micro-Tom genome assemblies reported previously as well as SL4.0 (Figure 1). This difference in length was caused by an approximately 15 Mb sequence at the top of the chromosome 2 (Figure 2), which was present in SLM_r2.0.pmol but absent in the other genome assemblies. It has been reported that an rDNA cluster is located at the top of chromosome 2 (Vallejos et al. 1986), and that rDNA clusters are commonly found in not only plant genomes but also animal genomes (Prokopowich et al. 2003). In tomato, to the best of our knowledge, no genome sequences for the rDNA cluster have been reported to date, since it might be difficult to sequence these repetitive DNA sequences with conventional NGS technologies, e.g., short-read and error-prone long-read methods. The high-fidelity long-read sequencing technology employed in this study was able to decode the sequence of complex genomic regions. The whole-genome sequencing analysis might provide new insights into the functions and evolutional history of genomes including complex structures.

Based on the chromosome-level genome assembly of Micro-Tom, EMS-induced mutations were detected in nine Micro-Tom mutant lines. The features of mutations in the nine lines, such as mutation density (1 mutation per 0.87 Mb), C/G to T/A transition rate (76.1%), and deleterious mutation rate (0.2%), were comparable with those reported in our previous study (Shirasawa et al. 2016b). In tomato, genes conferring fruit size have been well studied, including *fw2*.*2* (Frary et al. 2000), *locule number* (*lc*) (Muños et al. 2011), and *fasciated* (*fas*) (Cong et al. 2008). Indeed, *fas* has been suggested to control fruit size and shape in a miniature ornamental tomato cultivar (Safaei et al. 2020). In addition, in this study, 14 genes were selected as novel candidates that might enhance tomato fruit size. Even though further molecular genetic studies would be required to reveal the mechanisms controlling fruit size and development, a comprehensive list of gene candidates for large-fruit size in tomato could be obtained through whole-genome resequencing analysis, with the high-quality genome sequence serving as a reference.

In this study, we present a near-complete chromosome-level genome assembly of the Micro-Tom NBRP-Japan line, which showed the highest genome coverage compared with the previously reported genome assemblies of Micro-Tom and that of the standard line, Heinz 1706. The assembly revealed a structure at the top end of chromosome 2, where rDNA genes were predominantly clustered. The genome sequence data could become a new standard for functional genomics analysis of Micro-Tom, and might open new horizons for a detailed understanding of genomes including complex structures.

## Acknowledgments

We thank Y. Kishida, C. Minami, K. Ozawa, H. Tsuruoka, and A. Watanabe (Kazusa DNA Research Institute) for technical assistance. Seeds of Micro-Tom NBRP-Japan line (TOMJPF00001) and nine Micro-Tom mutant lines (TOMJPE2703, TOMJPE3484, TOMJPE5212-5, TOMJPE5406, TOMJPE5409, TOMJPE5770-1, TOMJPE5906, TOMJPW0604, and TOMJPW1559-1) were obtained from the University of Tsukuba, Tsukuba Plant Innovation Research Center, through the National Bio-Resource Project (NBRP) of MEXT/AMED, Japan.

## Data availability

Raw HiFi long reads of Micro-Tom NBRP-Japan line and short reads of Micro-Tom mutants were deposited in the Sequence Read Archive (SRA) database of the DNA Data Bank of Japan (DDBJ) under the accession numbers DRR503528–DRR503529 and DRR118572–DRR118580, respectively. The assembled sequences are available at DDBJ (accession numbers AP028935–AP028946) and KaTomicsDB (https://www.kazusa.or.jp/tomato).

## Author contributions

TA and KS conceived the project. TA prepared the plant materials. KS collected, analyzed, and interpreted the data. KS wrote the manuscript.

